# Subtle alteration in microRNA dynamics accounts for differential nature of cellular proliferation

**DOI:** 10.1101/214429

**Authors:** Dola Sengupta, Vinodhini Govindaraj, Sandip Kar

## Abstract

In the G_1_ phase of the mammalian cell cycle, a bi-stable steady state dynamics of the transcription factor E2F ensures that only a certain threshold level of the growth factor can induce a high expression level (on state) of E2F to initiate either normal or abnormal cellular proliferation or even apoptosis. A group of microRNA’s known as the mir-17-92 cluster, which specifically inhibits E2F, can simultaneously influence the threshold level of growth factor required for E2F activation, and the corresponding expression level of E2F in the on state. However, mir-17-92 cluster can function as either oncogene or tumor suppressor in a cell-type specific manner for reasons that still remain illusive. Here we put forward a deterministic mathematical model for Myc/E2F/mir-17-92 network that demonstrates how the experimentally observed mir-17-92 mediated differential nature of the cellular proliferation can be reconciled by having conflicting steady state dynamics of E2F for different cell types. While a 2-D bifurcation study of the model rationalizes the reason behind the contrasting E2F dynamics, an intuitive sensitivity analysis of the model parameters predicts that by exclusively altering the mir-17-92 related part of the network, it is possible to experimentally manipulate the cellular proliferation in a cell-type specific fashion for therapeutic intervention.

## Introduction

In mammalian cells, the expression levels of the key transcription factors E2F (mainly E2F1, E2F2 and E2F3) and Myc precisely govern the fate commitment of a mammalian cell toward either active cellular proliferation or quiescence under normal growth condition in the late G_1_ phase of the cell cycle [1–4]. Experimentally, it had been elegantly demonstrated that the expression level of the E2F1 protein follows a bi-stable dynamics (involving E2F1 and Myc) as a function of growth factor [1]. This implies that based on the expression levels of E2F1 in the late G_1_ phase, the mammalian cells can either opt for a quiescence state (Off state, low E2F1 expression) or can decide to go for the normal cell cycle (On state, high E2F1 expression). Fascinatingly, if the E2F1 protein level reaches very high expression level (On state) at relatively lower growth factor stimulation then the cells can even commit for apoptosis [1,2]. Whereas, an intermediate high expression level of E2F1 (between On state of normal cell cycle and apoptosis cases) maintained for an in-between threshold growth factor level can eventually lead to uncontrolled cellular proliferation or cancer [2]. Thus, finding out means to control the E2F1 dynamics in and around G_1_/S transition can be crucial from the therapeutic point of view.

In this regard, exploring the role of microRNA’s related to the mir-17-92 cluster seems worthwhile, as these microRNA’s are well-known antagonist of the E2F genes [5–10]. The mir-17-92 cluster consists of 7 different microRNA’s (mir-17-5p, mir-17-3p, mir-18a, mir-19a, mir-20a, mir-19b-1 and mir-92a-1), and is probably the most well studied microRNA cluster in the literature [5–10]. Several microRNAs associated with the mir-17-92 cluster play decisive roles in organizing the normal development as well as the events controlling the cellular proliferation in various cell types [11]. Like all the other microRNA’s, the members of the mir-17-92 cluster function post-transcriptionally by forming complexes with the mRNA’s of different target genes like E2F to either degrade or to reduce the translational efficiency of the corresponding target mRNA’s [2,12]. This seems to suggest that down regulating mir-17-92 cluster components may lead to unwanted cell cycle entry or cancer (as E2F levels will be high) at a lower threshold level of growth factor, which was indeed observed for many hematopoietic cancers [5,11,13,14]. Interestingly, for different solid type cancers, the microRNA’s of the mir-17-92 cluster were found to be mostly over-expressed [11,15–30]. This indicates that in solid cancer cells, the expression level of E2F must increase with the increasing level of microRNA’s related to mir-17-92 cluster. In this article, we want to rationalize this mir-17-92 cluster mediated differential dynamics of E2F protein in different cancer types, as it may provide us with crucial insights to prevent uncontrolled cellular proliferation and unwanted apoptosis events in a cell type dependent manner.

Disentangling the influence of mir-17-92 cluster on the cell type specific differential dynamics of E2F is not a straightforward problem. In fact, in mammalian cells, if we only focus on the mir-17-92 cluster mediated E2F1 dynamics, we will observe that it is tightly regulated by a number of feedback loops operative within a core network of Myc/E2F1/mir-17-92 [1,2,31–33] (Fig. 1). It is well known that E2F1 can positively regulate its own as well as Myc transcription [2,33]. Fig. 1 further depicts that both Myc and E2F1 can act as transcription factors for several microRNAs of mir-17-92 cluster, which in turn inhibit E2F1 [2,10,34–37]. This clearly suggests that in mammalian cells, either with or without mutation (or non-functional form) of Rb (Retinoblastoma protein, the main antagonist of E2F1), the cell fate decisions during G1-S transition can be modified by altering the expression level of mir-17-92 cluster related microRNAs. Still the question remains, how over expressing mir-17-92 cluster related microRNAs can fine-tune the dynamics of the Myc/E2F1/mir-17-92 network to generate differential activation dynamics of E2F1 in different cancer cell types?

**Fig. 1.**
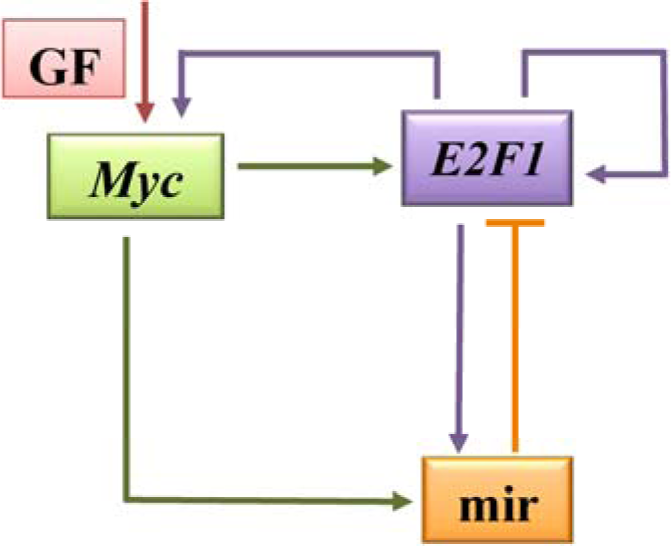
The growth factor regulated core interaction network of Myc/E2F1/mir-17-92. Solid arrows represent direct and indirect gene activation processes and the hammerhead from miR (representing mir-17-92 cluster) to E2F1 corresponds to the inhibition in the form of *E2F1* mRNA degradation or reduced level of translation.

In this article, we propose a mathematical model for the Myc/E2F1/mir-17-92 network illustrated in the Fig. 1 to resolve this important question. Our model reconciles the differential nature of the E2F1 dynamics under the overexpression condition of mir-17-92 cluster in a cancer cell type dependent fashion. The deterministic 2-dimensional bifurcation analysis of the model further substantiates why such differential E2F1 dynamics arises upon subtle alteration of the properties related to microRNAs of the mir-17-92 cluster. The model reproduces several experimental observations for cancer cell types of different origins as the expression level of mir-17-92 cluster is changed, and makes interesting experimentally testable predictions. We believe that our model essentially captures the circumstances under which mir-17-92 cluster behaves either as oncogene or as a tumor suppressor in a context dependent manner.

### The Myc/E2F1/mir-17-92 model

To develop our mathematical model, we have expanded the minimal gene regulatory network of Myc/E2F1/mir-17-92 shown in Fig. 1 by incorporating the crucial interactions based on existing experimental literature (Fig. 2). The model depicts that the growth factor (GF) activates the transcription of the Myc mRNA (*Myc*_m_) and *Myc*_m_ in turn produces Myc protein (Myc_p_). The Myc_p_ positively activates the transcription of both mir-17-92 cluster component (mir) and E2F1 mRNA (*E2F1*_m_). It is well established in the literature that E2F1 can only positively affect the transcription of its target genes if it forms a hetero-dimeric complex (DE) with the Dp protein (Dp1_p_) [31,33]. We have introduced this transcriptional positive feedback by the DE complex in the *Myc*_m_, mir and *E2F1*_m_ dynamics by using a phenomenological (Hill kinetic) term to represent the complex nature of the transcriptional activations in general. We have considered that both the Myc and E2F1 mRNAs and the corresponding proteins degrade with rates as observed in experiments [31,38–42]. Fig. 2 further describes that mir-17-92 cluster component (mir) provides a negative feedback to the E2F1 dynamics by forming a complex (Mpi) with the *E2F1*_m_. From this complex (Mpi) the *E2F1*_m_ gets degraded, and the translation efficiency of *E2F1*_m_ also gets reduced compare to its free form.

**Fig. 2.**
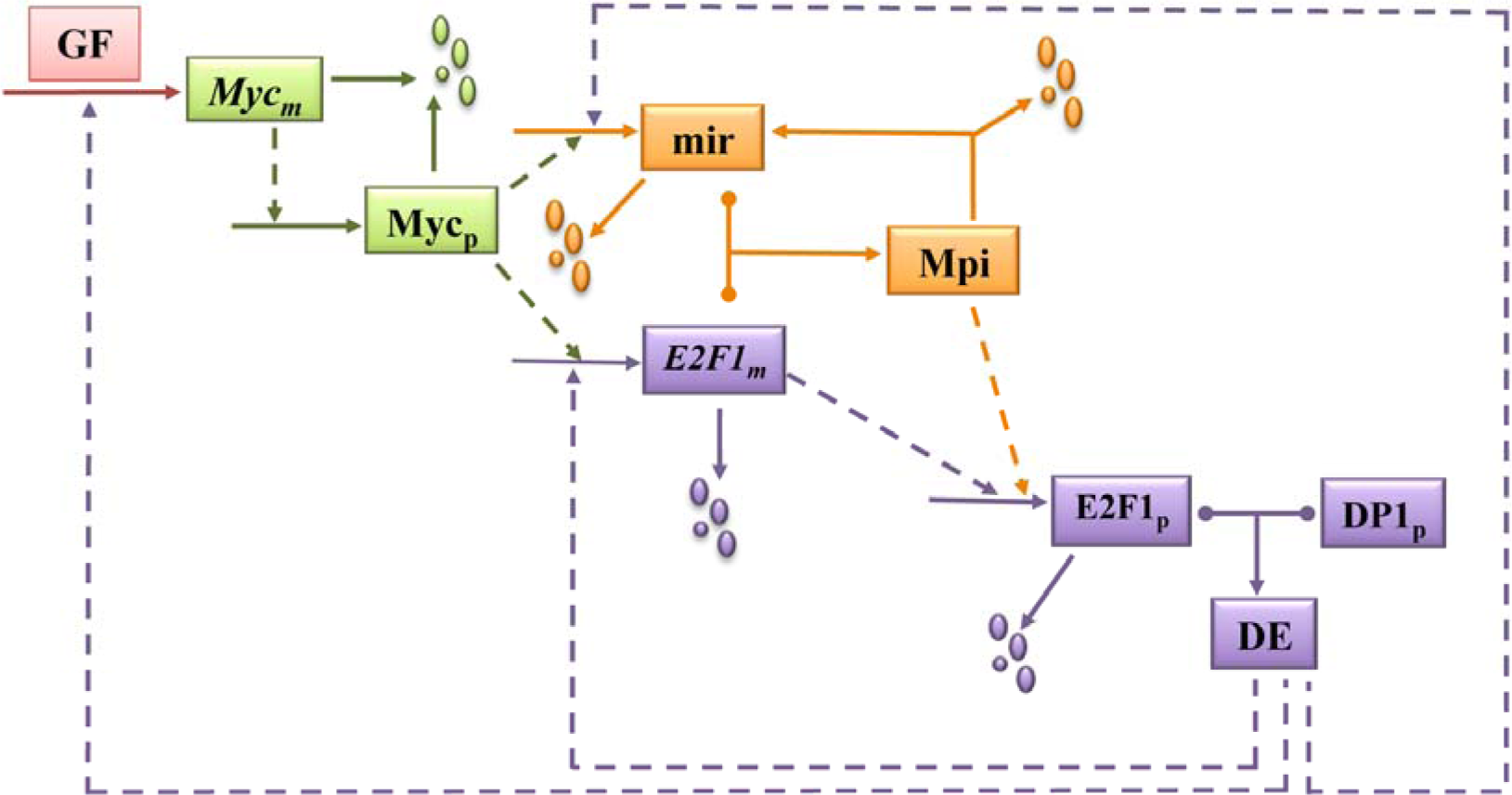
The Myc/E2F1/mir-17-92 regulatory network. (Solid and dashed arrows represent direct and indirect activation processes. Arrows pointing towards the dust (uneven circles) represents the process of mRNA, microRNA and protein degradation. Arrows coming out from the middle of a dumble shaped line indicates complexation and decomplexation events respectively.) More than one positive and negative feedback interactions are operative in this small but complex regulatory network.

We have translated the regulatory network shown in Fig. 2 into an ordinary differential equation based mathematical model (Table 1). The variables are defined in Table S1, and the definitions of the parameters along with their numerical values are provided in Table S2. We have acquired some of the parameter values directly from experimental literature [1,31,38–43] (shown in Table S2). It is worthwhile to mention that here we mainly focus to understand the emergent dynamic behavior of the regulatory network described in Fig. 2 to unravel the differential manner of E2F1 activation in cell-type specific manner as a function of varying expression level of mir-17-92 cluster component (mir). Thus, we have intuitively chosen the remaining parameters in the model in such a way that it allows us to explore the effect of different feedback loops present in the network by maintaining reasonable expression levels of all the mRNAs and proteins involved in the network.

**Table 1:**
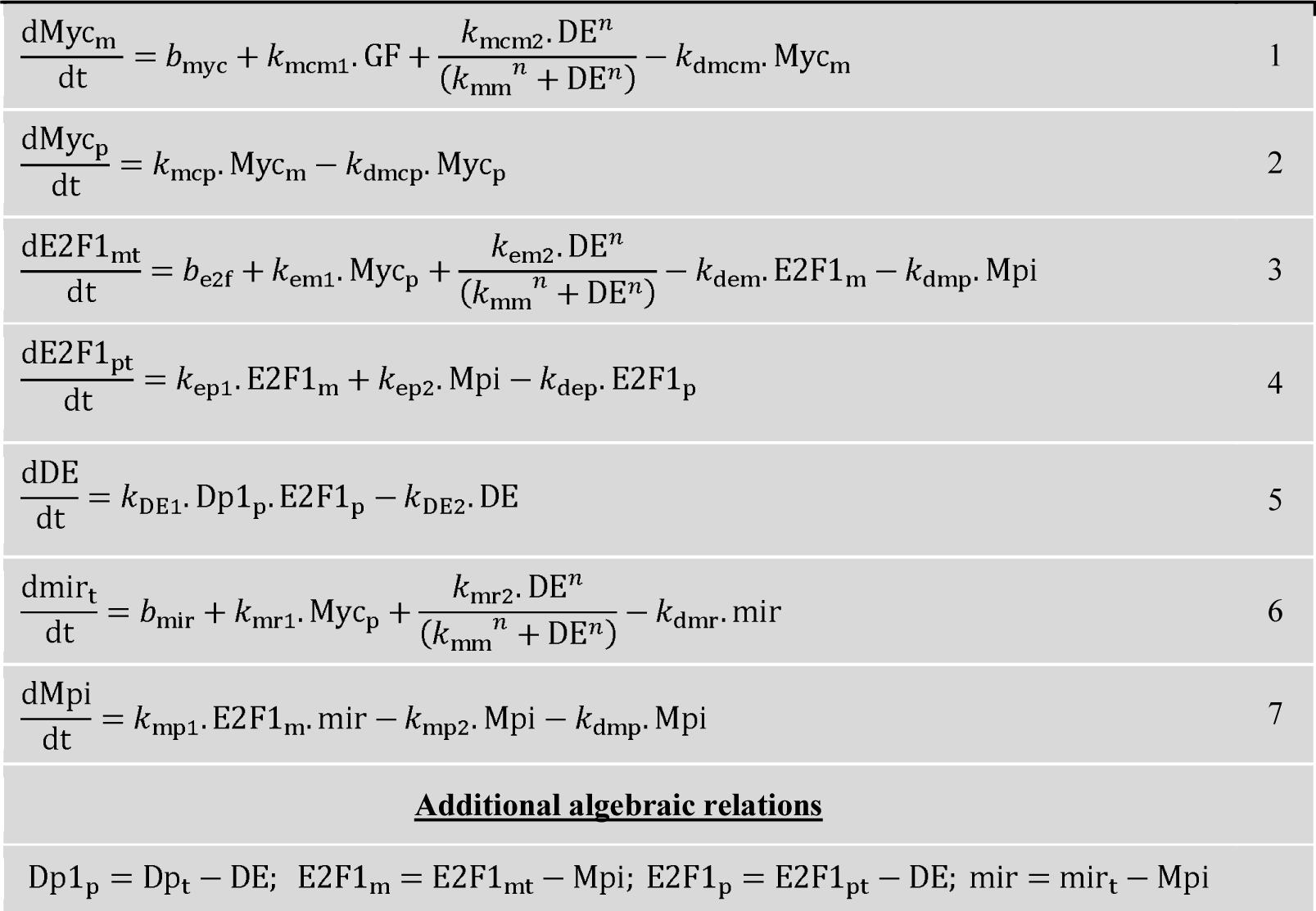
Equations governing the Myc/E2F1/mir-17-92 network

## Results

### E2F1 dynamics is indeed differentially regulated by mir-17-92 cluster components in a cell-type specific manner

To begin with, the bifurcation analysis of the model (Table 1) produces a typical bi-stable dynamics for the total E2F1 protein as a function of the growth factor (GF) that represents a wild type (WT, Fig. 3a) situation. The bi-stable dynamics ensures that the normal cell experiences a certain threshold of GF level to commit faithfully from the low level of E2F1 level (OFF state) to the higher E2F1 level (ON state) for the active proliferation cycle. An increase in the expression level of mir (situations (ii) and (iii) in Fig. 3a) not only increases the E2F1 level in the ON state, but makes the cells more susceptible for unwanted proliferation by simultaneously reducing (shifts the saddle node SN_1_ to the left of WT situation) the threshold requirement of GF. Expectedly, repression of mir (situation (i) in Fig. 3a) further increases the threshold requirement of GF (compared to WT). At this point, we fixed the GF level to 1 s.u., and showed (Fig. 3b) that for a cell type represented by a parameter set provided in Table S2, the steady state level of total E2F1 will increase with the increasing level of mir expression level. Fig. 3a-b resembles the kind of dynamics one observes in the cases of solid tumor cancer cells [11,15–30].

**Fig. 3.**
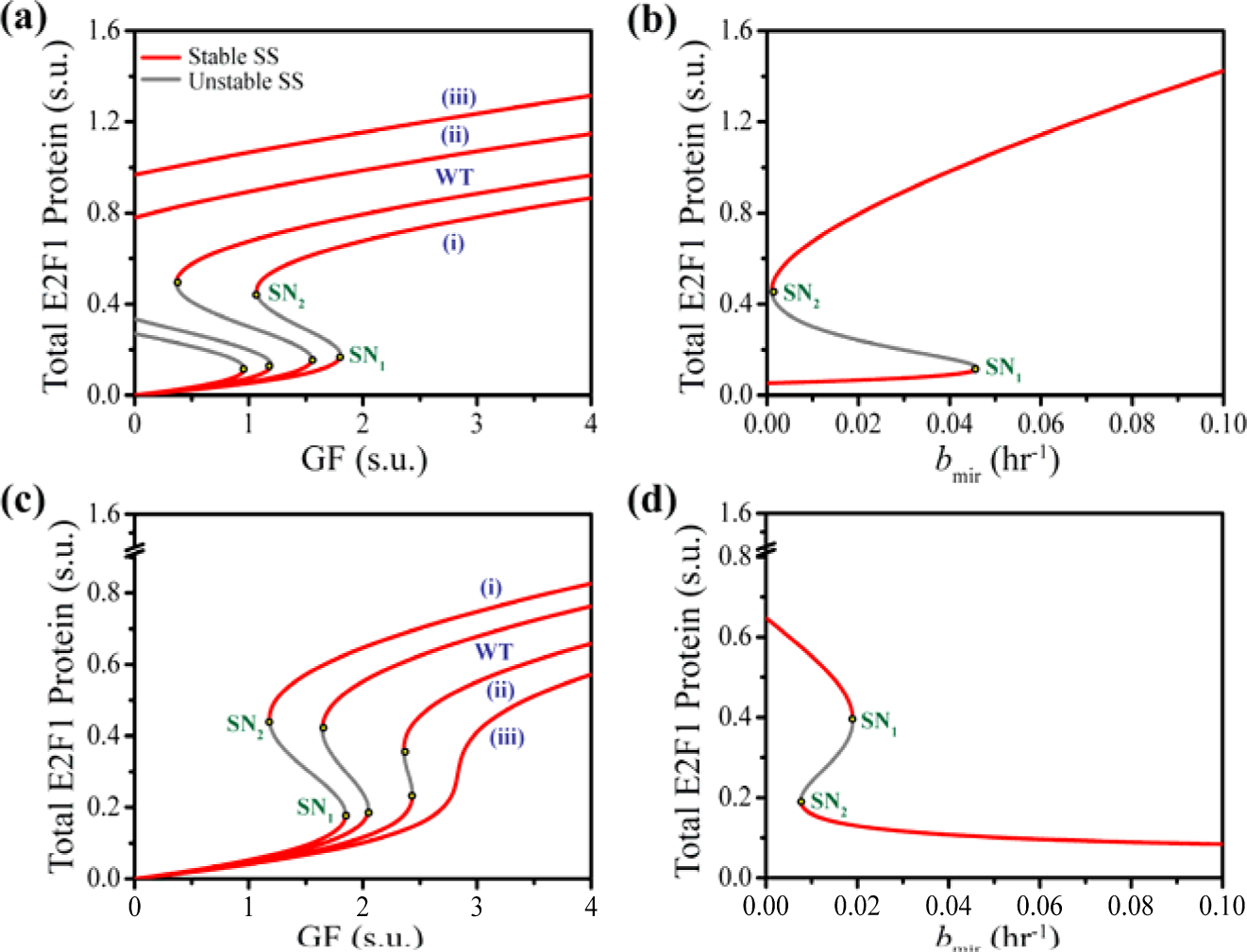
Bifurcation analysis of the model reveals the mir-17-92 mediated cell-type specific differential dynamics of E2F1. The steady state level of the total E2F1 protein (for the cell types where, ***increase*** in the mir expression leads to ***increase*** in the proliferation) is plotted as a functions of (**a**) growth factor (GF in scaled unit (s.u.)) for different basal activation rate (*b*_mir_) (0.01 hr^-1^ wild type (**WT**), 0.0001 hr^-1^ (**i**), 0.03 hr^-1^ (**ii**), 0.05 hr^-1^ (**iii**)) of the mir-17-92 cluster component (mir) and **(b)** basal activation rate (*b*_mir_) with a fixed level of growth factor (GF = 1 s.u.). Parameters values are given in Table S2. The steady state level of the total E2F1 protein (for the cell types where, ***increase*** in the mir expression leads to ***decrease*** in the proliferation) is plotted as a functions of **(c)** growth factor (GF in scaled unit (s.u.)) for different basal activation rate (*b*_mir_) (0.01 hr^-1^ wild type (**WT**), 0.0001 hr^-1^ (**i**), 0.03 hr^-1^ (**ii**), 0.05 hr^-1^ (**iii**)) of the mir-17-92 cluster component (mir) and **(d)** basal activation rate (*b*_mir_) with a fixed level of growth factor (GF = 2 s.u.). Parameters values are as given in Table S2, with only *k*_dmp_ = 0.15 hr^-1^ (c-d).

The question remains, how we can reverse the phenomenon keeping the same structure of the Myc/E2F1/mir-17-92 network. In this endeavor, we performed the bifurcation analysis after only changing the kinetic parameter *k*_dmp_ (from 0.01 hr^-1^ to 0.15 hr^-1^) that represents the rate at which mir facilitates the degradation of the *E2F1* mRNA from the Mpi complex. In Fig. 3c, the WT again represents the bi-stable switching E2F1 protein dynamics as a function of GF for a wild type situation of another specific cell type. Interestingly, an increase in the mir expression level now reduces the E2F1 steady state level (situations (ii) and (iii) in Fig. 3c) in the ON state by even creating a higher threshold value of GF for reaching the ON state as well. On the other hand, a decrease in the mir expression (Fig. 3c(i)) is elevating the propensity for proliferation by reducing the threshold requirement of GF. The bifurcation diagram (Fig.3d) of total E2F1 as a function of basal synthesis rate of mir further indicates that for a fixed GF level (2 s.u.), the increasing expression level of mir will decrease the E2F1 in this scenario. This fits the experimental observations found in some hematopoietic cancer cell types as well as for cells where, the mir-17-92 cluster components are found to act in an anti-apoptotic manner [5,11,13,14].

### The model validates the cell type dependent differential dynamics of E2F1

Next, we tried to validate our model outputs with the experimentally known phenotypes under two different scenarios discussed in the previous section. Cloonan et al. [44] have shown that only overexpressing a single micro-RNA (mir-17-5p) related to mir-17-92 cluster is sufficient to drive the HEK293T cells (solid tumor) into high level of proliferation. The MTT assay performed by them [44] revealed that the HEK293T cells became highly proliferative when mir-17-5p was stably overexpressed. At the same time, qRT-PCR experiment confirmed that the E2F1 expression level also increases significantly at the mRNA level [44].

Our model simulations (with *k*_dmp_ 0.01 hr^-1^) predict that in comparison to the WT case, overexpressing the mir-17-92 component elevates the level of *E2F1* mRNA (Fig. 4a, left panel) up to an extent observed in the experiments [44]. Interestingly, the temporal evolution dynamics of E2F1 protein (Fig. 4a, right panel) under mir-17-92 overexpression condition seems to correlate nicely with proliferation profile observed in the MTT assay performed by Cloonan et al. [44].

**Fig. 4.**
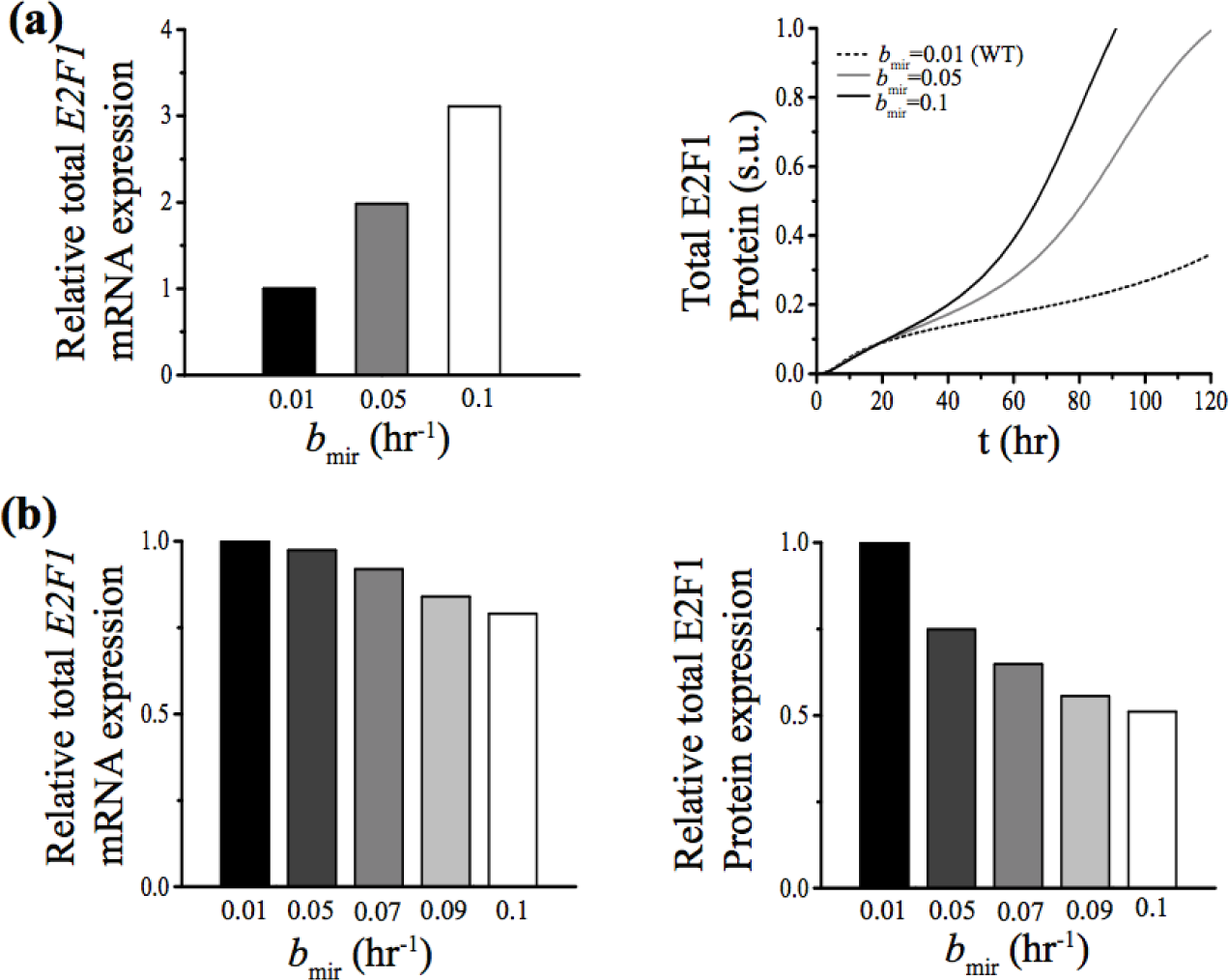
Overexpression of mir-17-92 cluster components reproduce experimentally observed differential activation of E2F1 protein in cell-type specific manner. **(a)** Relative expression levels of total *E2F1* mRNA (left panel) as a function of increased level of mir (at GF=2 s.u. and other parameters as in Table S2). Time profiles for total E2F1 protein level that are plotted as a function of time (right panel) for different values of basal activation rate of mir (*b*_mir_) correlate with the increase in the rate of proliferation when the mir-17-92 cluster components are overexpressed in HEK293T cells. **(b)** The model predicted relative expression levels of total *E2F1* mRNA (left panel) and protein (right panel) as a function of increased level of mir corroborates with experimentally observed levels of E2F1 protein and mRNA in Hela cells (at GF=4 s.u., *k*_dmp_ = 0.15 hr^-1^and other parameters as in Table S2).

On the other hand, it has been observed experimentally in case of Hela cells [35], transient overexpression of the mir-17 cluster has little or no effect on the E2F1 mRNA expression, but it decreases the E2F1 protein expression by about 50%. Our model simulations (with *k*_dmp_ 0.15 hr^-1^) suggest that the E2F1 protein level can be reduced to 50% of that of the WT situation by overexpressing the mir-17-92 level (Fig. 4b, right panel). Under the same condition the mRNA level of the *E2F1* shows very little effect (Fig. 4b, left panel). This further emphasizes that mir-17-92 can either enhance or reduce the extent of proliferation by just differentially altering the E2F1 dynamics in a cell type dependent fashion.

### Two-parameter bifurcation analysis unravels the dynamical origin of differential dynamics of E2F1

In Fig. 3 and Fig. 4, we have shown that just by altering *k*_dmp_ i.e., the microRNA mediated degradation rate of *E2F1* mRNA, it is possible to rationalize the differential E2F1 dynamics. To understand how the changes in the parameter *k*_dmp_ affects such an interesting dynamical transition, we performed a two-parameter bifurcation analysis of the model (Fig. 5) by setting different expression level of the mir-17-92. In Fig. 5, we have overlaid the two-parameter bifurcation diagrams (*k*_dmp_ Vs GF) or the cusps obtained for the overexpressed (red line) and down-regulated (green line) mir-17-92 situations on top of the cusp (blue line) obtained for the WT expression level of mir-17-92. Interestingly, all the three cusps intersects at *k*_dmp_ ~ 0.083 hr^-1^ as indicated by the yellow dots in Fig. 5. This evidently clarifies why our chosen values of *k*_dmp_ (i.e., 0.01 hr^-1^ and 0.15 hr^-1^, indicated by red dots) allow us to reconcile the cell-type dependent differential E2F1 dynamics. Fig. 5 indicates that for *k*_dmp_ = 0.01 hr^-1^ if one goes on increasing the GF level, the overexpressed mir-17-92 situation (red line) appears before the WT situation (blue line), which follows the mir-17-92 down-regulated case (green line).

**Fig. 5.**
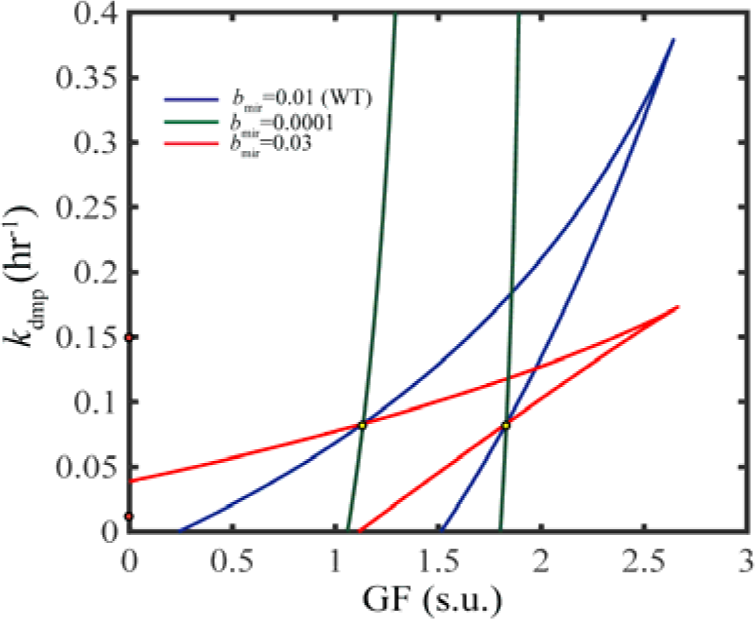
The 2-D bifurcations provide dynamical insight about the cell-type dependent differential dynamics of E2F1. Two-parameter (*k*_dmp_ Vs GF) bifurcation diagram is plotted (based on steady state levels of total E2F1 protein) for three different values (0.01 hr^-1^ (blue line), 0.0001 hr^-1^ (green line), 0.03 hr^-1^ (red line)) of basal activation rate of mir. The orange spots on *k*_dmp_-axis show the values of *k*_dmp_ used in Fig 3a and 3c. The yellow dots are the points where, the three two-parameter diagrams intersect in the *k*_dmp_ -GF plane and hence provide the critical value of *k*_dmp_.

Thus, it qualitatively shows why for *k*_dmp_ = 0.01 hr^-1^ situation if we overexpress mir-17-92, the cells transit to the S-G_2_-M phase (On state) earlier compare to the WT (observed in solid tumor [11,15–30]). The scenario reverses for the case with *k*_dmp_ = 0.15 hr^-1^, as now with increasing level of GF, the mir-17-92 down-regulated phenotype or the green line will appear even before the WT case (blue line). Consequently, the mir-17-92 down-regulated phenotype will commit to proliferation even at lower level of GF compare to the WT (as observed in some cases of hematopoietic cancer [5,11,13,14]).

We have shown further that even by altering the translational efficiency (*k*_ep2_) of the mir-17-92 bound E2F1 mRNA (Fig. S1), the cell type dependent E2F1 dynamics can also be reconciled. The important question one can ask is whether it is possible to manipulate either *k*_dmp_ or *k*_ep2_ experimentally to influence the cell fate decision-making process or not? In our model, the way we modeled the microRNA and *E2F1* mRNA interaction is quite simplified in nature. In literature it is already established that microRNAs need the presence of Argonaute proteins (especially Ago2 protein in mammalian cells) to either degrade mRNAs following different degradation pathways or even reducing the translational efficiency of the mRNAs by localizing them into P-bodies [45]. Thus, by either overexpressing or knocking down the Ago2 protein depending on the cell type, we can influence the commitment towards proliferation by altering the E2F1 dynamics.

### Myc gene related phenotypes could be reconciled for different levels of mir-17-92

Our minimal model of Myc/E2F1/mir-17-92 network is constructed in such a way that it allows us to investigate the importance of the Myc in the network. Experimentally it has been observed that for conditional lymphoma and leukemic cell lines, overexpressing the microRNAs from mir-17-92 cluster can rescue the cells where, *Myc* gene expression is temporarily inactivated upon doxycycline treatment [46]. Our model simulation (Fig. 6a, upper panel) shows that once the Myc gene expression is switched off, the relative E2F1 protein expression level falls off very rapidly in 3-4 days time as observed in the experiments [46] for the control cells kept under a basal GF medium. Intriguingly, if the mir-17-92 is overexpressed in the model as soon as the Myc gene is repressed, the relative E2F1 protein level seems to increase gradually (Fig. 6a, lower panel) with time corroborating the experimental findings [46].

**Fig. 6.**
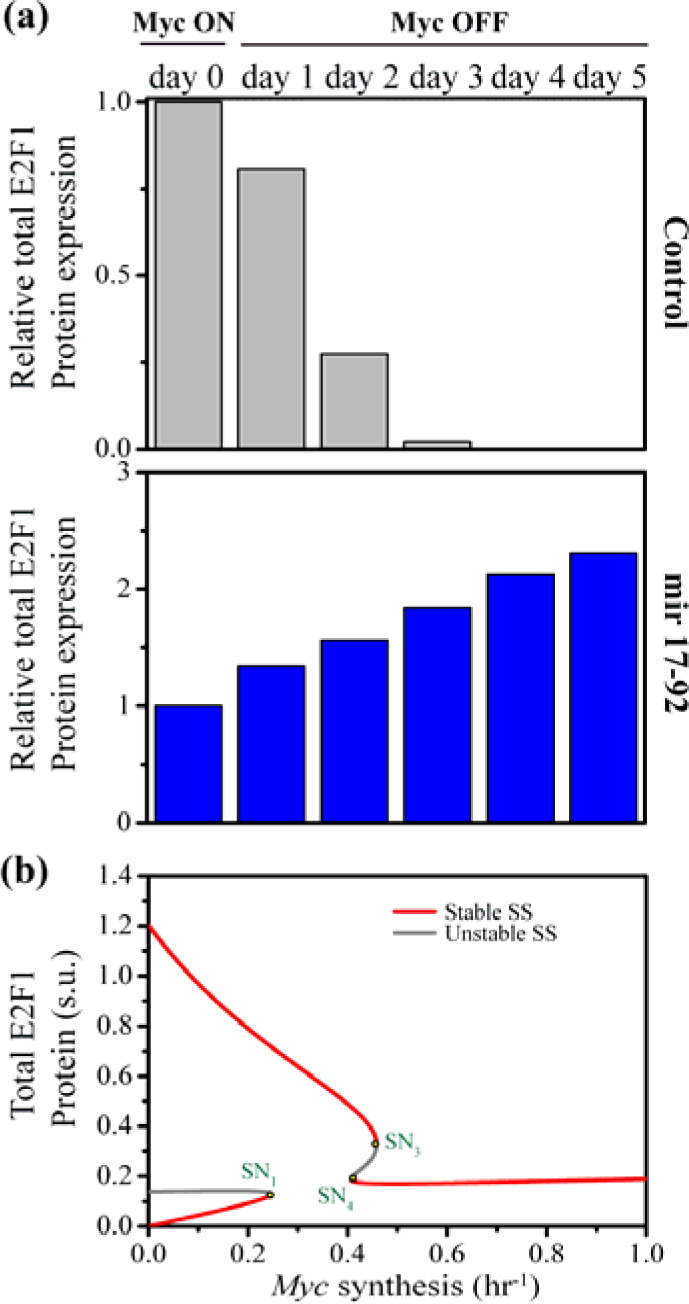
Model predictions related to Myc agree well with the experimental observations. **(a)** In control cells, the relative total E2F1 protein expression goes down quite rapidly with time if the *Myc* expression is switched off (Upper panel). Whereas, in mir overexpressed cells under the same situation, the E2F1 expression level increases progressively with time (lower panel) as observed in experiments. **(b)** The bifurcation diagram of the total E2F1 protein as a function of basal *Myc* synthesis rate (*b*_myc_) for a specific cell type (REF52 cells) under a basal growth factor (GF=0.0001 s.u.) condition. For very low and relatively high values of *b*_myc_, the expression level of E2F1 is low but for a in between *b*_myc_ level the E2F1 expression is relatively high which is in line with experimental observation. For details see the method section.

Our model further describes a Myc phenotype, which is even more challenging to envisage. It has been displayed experimentally [42] that overexpressing *Myc* gene systematically in serum starved REF52 cells containing a d2GFP reporter driven by E2F1 promoter produces a unique phenotype where, for very low and very high level of *Myc* gene expression, one can have relatively low level of E2F1 protein expression, but a moderate level of *Myc* expression leads to high expression level of E2F1. The steady state analysis of our model suggests that for such serum starved cells (kept under GF = 0.0001 s.u.), the E2F1 steady state will produce a mushroom like bifurcation (Fig. 6b) as a function of the basal synthesis rate of the *Myc*.

This evidently shows why for very low and very high expressions of *Myc* mRNA, a low steady state level of E2F1 expression can be achieved, while a moderate level of *Myc* will lead the dynamical system to the higher expressing E2F1 steady state.

### Sensitivity analysis reveals ways to influence the cell-type dependent proliferation response

At this juncture, we performed sensitivity analysis (Figs. S2 and S3 for *k*_dmp_ = 0.01 hr^-1^, and Figs. S4 and S5 for *k*_dmp_ = 0.15 hr^-1^) of the model parameters by taking the positions of SN_1_ and SN_2_ as the sensitivity parameter to explore the possible ways to influence the cell-type dependent proliferation by adjusting the E2F1 dynamics.

A careful inspection of the sensitivity diagrams (Figs. S2-S5) allowed us to separate out the parameters (as summarized in Fig. 7a-b), which would have differential impact on the respective saddle nodes for the two chosen values of *k*_dmp_. Interestingly, all the parameters that appear in Fig. 7, in one way or other are related to the mir-17-92 cluster component, and one of them is indeed the *b*_mir_. Fig. 7 opens up a set of experimental possibilities to change the fate of a proliferating cell just by manipulating the E2F1 dynamics. For example, depending on the requirement, (i) the *k*_dmr_ can be altered by producing a more or less stable variant of microRNA, (ii) *k*_mp1_ can be varied experimentally to reduce the association propensity of mir and mRNA of E2F1 and (iii) *k*_mr1_ and *k*_mr2_ can be manipulated by blocking the transcription initiation sites or reducing the level of other transcriptional co-activator require for mir production. Thus, the model qualitatively predicts a set of selective routes to control the proliferation by just perturbing the Myc/E2F1/mir-17-92 network.

**Fig. 7.**
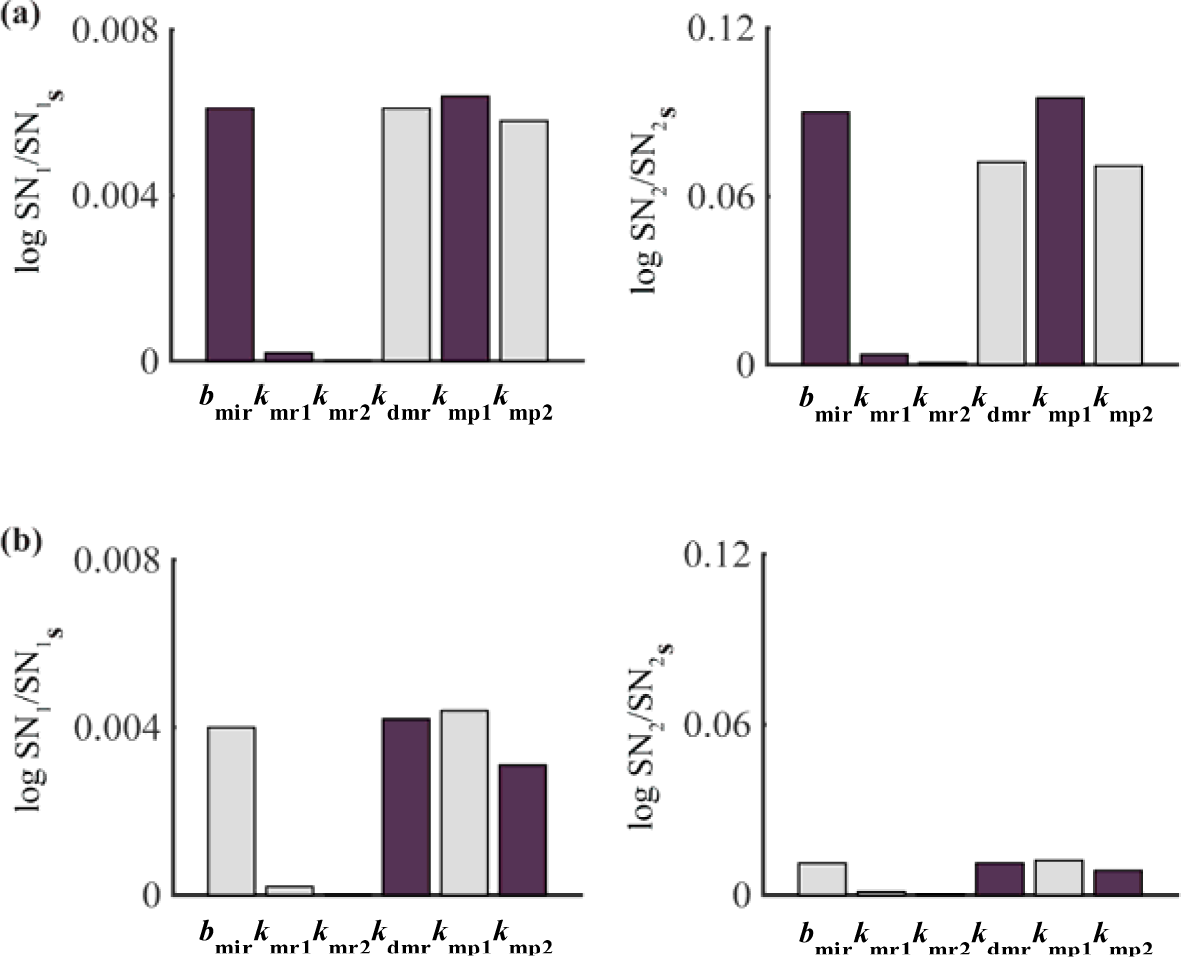
Sensitivity analysis provides crucial leads to alter cellular proliferation in a cell-type dependent manner. Sensitivity analysis of the model parameters is performed by taking the positions of SN_1_ and SN_2_ as the sensitivity parameter for the WT bifurcation diagrams drawn (as shown in Fig. 3a and c) for **(a)** *k*_dmp_ = 0.01 hr^-1^ and **(b)** *k*_dmp_ = 0.15 hr^-1^. (The different color bars signify that the corresponding saddle node will move towards either lower (purple bar) or higher (light grey bar) GF with respect to the WT scenario, if a particular parameter involved in the model is increased individually by about 10% of its original value (Table S2) keeping all other parameters constant.)

## Discussion and Conclusions

In many cancerous cells, the principle inhibitor of E2F1 i.e., the Retinoblastoma protein (Rb) remains either as mutated or non-functional [17]. Under such a condition, the microRNAs related to mir-17-92 can play decisive role in directing the cells toward either cellular proliferation or apoptosis or even in some cases to quiescence by controlling the E2F1 dynamics. Thus, mir-17-92 can act as oncogene as well as tumor suppressor depending on the cell type [2,5,11,13–30]. To understand the intricacies of the interactions between E2F1, Myc and microRNAs related to mir-17-92 cluster, we proposed and investigated a relatively simple Myc/E2F1/mir-17-92 network (Fig. 2). Our model essentially captures all the positive and negative feedback interactions that are responsible for the bi-stable E2F1 dynamics observed in mammalian cells. The 1-D bifurcation analysis (Fig. 3) of our model qualitatively depicts the experimentally observed differential dynamics of E2F1 protein (as a function of GF) under different mir expression levels. This not only corroborates (Fig. 4) with the experimental findings for different solid cancers, but also supports the experiments, where overexpressing mir-20a and mir-17-5p caused increased level of apoptosis [11,15–30]. In the latter case the experiments further suggest that the apoptosis is caused due to increased expression of E2F1 protein upon down regulating the mir-20a and mir-17-5p [34].

Importantly, the 2D-bifurcation diagrams (Fig. 5 and Fig. S1(b)) reveal the one of the probable routes by which the reversal of the E2F1 steady state dynamics can be achieved under overexpressing condition of mir, and how it can be realized by just altering the *k*_dmp_ or *k*_ep2_. Intriguingly, the sensitivity analysis (Fig. 7) of the model uncovers that the reversal of the E2F1 dynamics can only be accomplished by changing the mir-17-92 related parameters. We further discussed how one could perform experiments at different levels to test these predictions. In this context, it is worthwhile mentioning that our model definitely improves the current understanding about the Myc/E2F1/mir-17-92 network than the previous model proposed by Aguada et al. [2]. Although their model was a simpler one compare to us, but it failed to portray how the steady state behavior of E2F1 protein can vary in an opposite manner (as a function of GF) as the mir is overexpressed in a cell type dependent manner. Not only that, our model has a broader applicability, as it also adequately explains the Myc related mutants (Fig. 6) under different expression levels of mir [42,46].

To conclude, our proposed model effectively replicates the mir-17-92 influenced steady state dynamics of E2F1 in a cell-type dependent manner. It predicts exactly how one can manipulate the mir related part of the Myc/E2F1/mir-17-92 network to push a particular cancerous cell to either quiescence or apoptosis. Thus, it provides important insights to utilize the oncogenic or tumor suppressor behavior of mir-17-92 cluster in a context dependent manner for developing novel therapeutic strategies to treat uncontrolled proliferation.

## Methods

### Deterministic analysis

The complete gene regulatory network (Fig. 1) was constructed in terms of 7 ordinary differential equations. The deterministic bifurcation analysis of the model was executed using the freely available software XPP-AUT. The bifurcation diagrams and time profiles were drawn in Origin Lab using the data points generated by XPP-AUT.

In the systematic sensitivity analysis section, each parameter was increased individually at an amount of 10% with respect to the values given in Table S2 keeping all other parameters constant (Figs. S2, S3, S4 and S5). Bar diagrams were drawn with the help of MATLAB. Purple colored bar depicts the movement of the saddle nodes (SN_1_ and SN_2_) towards lower GF level and light-grey colored bar signifies movement of the saddle nodes (SN_1_ and SN_2_) towards higher GF than the WT cell-type specific cases.

In Fig. 6(a) to reconcile Myc ON state we have used *b*_myc_ = 0.5 hr^-1^ and *b*_myc_ = 0 hr^-1^ represents Myc OFF state and the simulations are performed at low GF (GF=0.0001 s.u.) and in absence of the positive feedback of E2F1 on Myc (*k*_mcm2_ = 0 hr^-1^). Other parameters are same as in Table S2. To draw the bifurcation diagram of total E2F1 protein as a function of *b*_myc_ (Fig. 6b) *k*_ep2_ = 0.014 hr^-1^, *k*_em2_ = 0.03 hr^-1^, *k*_mr1_ = 0.9 hr^-1^and GF = 0.0001 s.u. are used.

## Acknowledgements

Thanks are due to IRCC, IIT Bombay (13IRTAPSG005) for a fellowship to (DS). This work is supported by the funding agencies IRCC, IIT Bombay (13IRCCSG008), DST grant (EMR/2014/000500) and DBT grant (BT/PR11932/BRB/10/1315/2014).

## Conflict of Interest

The authors declare that they have no conflict of interest.

